# Building Large Updatable Colored de Bruijn Graphs via Merging

**DOI:** 10.1101/229641

**Authors:** Martin D. Muggli, Bahar Alipanahi, Christina Boucher

## Abstract

**Motivation:** There exists several massive genomic and metagenomic data collection efforts, including GenomeTrakr and MetaSub, which are routinely updated with new data. To analyze such datasets, memory-efficient methods to construct and store the colored de Bruijn graph have been developed. Yet, a problem that has not been considered is constructing the colored de Bruijn graph in a scalable manner that allows new data to be added without reconstruction. This problem is important for large public datasets as scalability is needed but also the ability to update the construction is also needed.

**Results:** We create a method for constructing and updating the colored de Bruijn graph on a very-large dataset through partitioning the data into smaller subsets, building the colored de Bruijn graph using a FM-index based representation, and succinctly merging these representations to build a single graph. The last step, merging succinctly, is the algorithmic challenge which we solve in this paper. We refer to the resulting method as VariMerge. We validate our approach, and show it produces a three-fold reduction in working space when constructing a colored de Bruijn graph for 8,000 strains. Lastly, we compare VariMerge to other competing methods — including Vari (Muggli *et al*., 2017), Rainbowfish (Almodaresi *et al*., 2017), Mantis (Pandey *et al*., 2018), Bloom Filter Trie (Holley *et al*., 2016), the method by Almodaresi *et al*. (2019) and Multi-BRWT (Karasikov *et al*., 2019)—and illustrate that VariMerge is the only method that is capable of building the colored de Bruijn graph for 16,000 strains in a manner that allows additional samples to be added. Competing methods either did not scale to this large of a dataset or cannot allow for additions without reconstruction.

**Availability:** Our software is available under GPLv3 at https://github.com/cosmo-team/cosmo/tree/VARI-merge.

**Contact:** Martin D. Muggli

## 1 Introduction

The money and time needed to sequence a genome has decreased remarkably in the past decade. With this decrease has come an increase in the number and rate at which sequence data is collected for public sequencing projects. This led to the existence of GenomeTrakr, which is a large public effort to use genome sequencing for surveillance and detection of outbreaks of foodborne illnesses. This effort includes over 50,000 samples, spanning several species available through this initiative—a number that continues to rise as datasets are continually added (Stevens *et al*., 2017). Another example is illustrated by the sequencing of the human genome. The 1000 Genomes Project (The 1000 Genomes Project Consortium, 2015) was announced in 2008 and completed in 2015, and now the 100,000 Genomes Project is well underway (Turnbull *et al*., 2018). Unfortunately, methods to analyze these and other large public datasets are limited due to their size.

Iqbal *et al*. (2012) presented one method for analysis sequence data from a large population which focuses on the construction of the *colored de Bruijn graph*. To define the colored de Bruijn graph, we first define the traditional de Bruijn graph and then show how it can be extended. Formally, a de Bruijn graph constructed for a set of strings (e.g., sequence reads) has a distinct vertex *v* for every unique (*k* − 1)-mer (substring of length *k* − 1) present in the strings, and a directed edge (*u, v*) for every observed *k*-mer in the strings with (*k* − 1)-mer prefix *u* and (*k* − 1)-mer suffix *v*. In the colored de Bruijn graph, the edge structure is the same as the classic structure, but now to each node ((*k* − 1)-mer) and edge (*k*-mer) is associated a list of colors corresponding to the samples in which the node or edge label exists. More specifically, given a set of *n* samples, there exists a set 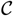 of *n* colors *c*_1_, *c*_2_,…, *c_n_* where *c_i_* corresponds to sample *i* and all *k*-mers and (*k* − 1)-mers that are contained in sample *i* are colored with *c_i_*. A *bubble* in this graph corresponds to a directed cycle, and is shown to be indicative of biological variation. Iqbal *et al*. (2012) show after construction of the colored de Bruijn graph, they can recover all genetic variants in the underlying population by traversing the graph, and finding paths with bubbles (“bubble finding”).

A bottleneck in applying Cortex (Iqbal *et al*., 2012) to large datasets lies the amount of memory and disk space required to build and store the colored de Bruijn graph. Thus, in recent years, there has been an initiative to build and/or store the colored de Bruijn graphs in a space-and time-efficient manner allowing it to be applied to larger populations. Vari (Muggli *et al*., 2017), Rainbowfish (Almodaresi *et al*., 2017), Mantis (Almodaresi *et al*., 2019), the method of Karasikov *et al*. (2019), as well as others, sought to overcome this limitation by improving the storage efficiency of the graph. Vari was one of the first methods to build the colored de Bruijn graph in a memory efficient manner. It extends the de Bruijn graph construction of Bowe *et al*. (2012). Rainbowfish (Almodaresi *et al*., 2017) was later developed. It first uses Vari to build the colored de Bruijn graph, which it compresses further by storing identical rows in the Vari color matrix as a single row. Cortex, Vari and Rainbowfish have bubble-calling methods that allow genetic variation between the datasets to be detected.

Mantis (Almodaresi *et al*., 2019) improve on Vari and Rainbowfish by constructing a different data structure, which it is able to construct directly without using Vari. Most recently, the method of Almodaresi *et al*. (2019) was presented, which is referred to as MST-based color-class representation. This method first uses Mantis to build the colored de Bruijn graph, which it then compresses further using minimum spanning trees and storing only deltas between similar vectors. Both Mantis and the method of Almodaresi *et al*. (2019) do not have bubble-calling procedures. We note there exists more compressed colored de Bruijn graph methods, and give a comprehensive review in the next section.

Unfortunately, there does not exist a method to build the colored de Bruijn graph in a manner that is scalable to large datasets and that allows for the addition of new data. This is important since one of the original purposes of the colored de Bruijn graph was to analyze population-level datasets — such as GenomeTrakr and 100,000 Genomes Project — which continually grow in size due to the addition of new data. Existing colored de Bruijn graph methods either cannot scale well enough to construct for large datasets, or cannot update the data structure without complete reconstruction. For example, Vari (Muggli *et al*., 2017), Rainbowfish (Almodaresi *et al*., 2017), Mantis (Almodaresi *et al*., 2019) and the method Karasikov *et al*. (2019) are efficient with respect to memory and time but cannot be updated without reconstructing the entire colored de Bruijn graph. Bloom Filter Trie (Holley *et al*., 2016) and the method of Karasikov *et al*. (2019) allow for the addition of new data but cannot scale to large datasets, as we show in this paper. Crawford *et al*. (2019) build a de Bruijn graph that can be updated but restrict interest to the traditional (non-colored) de Bruijn graph.

Here, we focus on scalable construction of the colored de Bruijn graph in a manner that allows new data to be added without reconstruction. One way to achieve this construction is to devise a divide-and-conquer approach which will divide the complete set of samples (e.g. sets of sequence reads) into smaller sets, construct a succinct colored de Bruijn graph for each smaller set, and merge these succinct colored de Bruijn graphs into progressively larger graphs until a single one remains. The problem of merging succinct colored de Bruijn graphs efficiently is a challenging problem but necessary to solve since it avoids memory, disk and time overhead. This divide-and-conquer approach also allows the graph to be updated. Given new sets of sequence reads, they can be used to build a compressed colored de Bruijn graph and then merged into the existing (larger) compressed colored de Bruijn graph.

### Our contributions

We present VariMerge which constructs a colored de Bruijn graph for large datasets in a manner that allows new datasets to be added efficiently. As suggested earlier, VariMerge builds through a process of partitioning the data into smaller sets, building the colored de Bruijn graph in a RAM-efficient manner for each partition, and merging colored de Bruijn graphs. Each of the colored de Bruijn graphs is stored using the FM-index in the same manner as Vari (Muggli *et al*., 2017). Thus, the algorithmic challenge that we tackle is merging the graphs in a manner that keeps them in their compressed format throughout the merging process — rather than decompressing, merging and compressing, which would be impractical with respect to disk and memory usage.

We verify the correctness of our approach and the accompanied bubble-calling algorithm by showing the colored de Bruijn graph built via merging is bit-for-bit identical to standard construction, and successfully identifies all bubbles in the merged graph. Next, we demonstrate that VariMerge improves construction scalability over Vari, reducing running time by a third and a reducing working space three fold by comparing the peak disk and memory required to build the colored de Bruijn graph for 8,000 Salmonella strains using Vari and VariMerge.

Lastly, we use VariMerge to build a colored de Bruijn graph for 16,000 strains of Salmonella. This construction required 254 GB of RAM, 2.34 TB of external memory, and approximately 70 hours of CPU time. To contextualize these results we compare the construction of VariMerge with those of state-of-the-art methods, including Vari (Muggli *et al*., 2017) / Rainbowfish (Almodaresi *et al*., 2017)^1^, Bloom Filter Trie (Holley *et al*., 2016), Mantis (Pandey *et al*., 2018) / the method of Almodaresi *et al*. (2019)^2^, and the method of Karasikov *et al*. (2019), on datasets consisting of 4,000, 8,000 and 16,000 strains. We demonstrate the incremental update utility of VariMerge by adding a new strain to the 16,000 set an order of magnitude faster that its initial construction.

These results demonstrate that VariMerge is the only method that is capable of building the colored de Bruijn graph for 16,000 strains in a manner where new datasets can be added (via merging). Mantis was the only other method that was capable of building the colored de Bruijn graph for 16,000 strains. Although Mantis (Pandey *et al*., 2018) was able to constructed a colored graph for 16,000 strains, and the authors suggest a dynamic update strategy in future work, no implementation is available and its current implementation cannot be updated without reconstruction. All other methods were not scalable and thus, we were unable to complete construction in 72 hours and using at most 4 TB of disk space and 756 GB of memory.

## 2 Related Work

### Efficient de Bruijn graph

Space-efficient representations of de Bruijn graphs have been heavily researched in recent years. One of the first approaches was introduced with the creation of the ABySS assembler, which stores the graph as a distributed hash table (Simpson *et al*., 2009). (Conway and Bromage, 2011) reduced these space requirements by using a sparse bitvector by (Okanohara and Sadakane, 2007) to represent the *k*-mers, and used rank and select operations (to be described shortly) to traverse it. Minia (Chikhi and Rizk, 2013) uses a Bloom filter to store edges, which requires the graph to be traversed by generating all possible outgoing edges at each node and testing their membership in the Bloom filter. (Bowe *et al*., 2012) developed a succinct data structure based on the Burrows-Wheeler transform. This data structure is discussed in more detail in the next section. This data structure of (Bowe *et al*., 2012) is combined with ideas from IDBA-UD (Peng *et al*., 2012) in a metagenomics assembler called MEGAHIT (Li *et al*., 2015). (Chikhi *et al*., 2014) implemented the de Bruijn graph using an FM-index and minimizers.

### Efficient colored de Bruijn graphs

As previously mentioned, Vari (Muggli *et al*., 2017) and Rainbowfish (Almodaresi *et al*., 2017) are both space efficient data structures for storing the colored de Bruijn graph which use the structure of (Bowe *et al*., 2012). (Muggli *et al*., 2017) build the compressed color matrix by compressing each row using Elias-Fano encoding. Rainbowfish (Almodaresi *et al*., 2017) takes as input the color matrix of Vari and compress it by decomposing the matrix into “color sets” based on an equivalence relation and compresses each color set individually. Thus, this method relies on the construction of Vari prior to building the sets of compatible colors. (Holley *et al*., 2016) introduced the Bloom Filter Trie, which is another succinct data structure for the colored de Bruijn graph. It encodes frequently occurring sets of colors separate from the graph and stores a reference to the set if the reference takes fewer bits than the set itself. This data structure allows incremental updates of the underlying graph. More recently, Mantis (Pandey *et al*., 2018) and its extension (Almodaresi *et al*., 2019) improve upon Vari and Rainbowfish. They build a compressed colored de Bruijn graph by building sets of compatible colors (similar to Rainbowfish) but construct the compressed graph directly rather than constructing it from Vari. Lastly, the most recent method (Almodaresi *et al*., 2019), constructs the colored de Bruijn graph of Mantis and then further compresses the color matrix constructed from Mantis.

### Compression of Color Matrix

There have been a couple methods that focus on constructing the color matrix in a manner that is both compressed and/or dynamic. This includes the method of Mustafa *et al*. (2018) and Multi-BRWT (Karasikov *et al*., 2019). The main contribution of these methods is that they build the color matrix in a manner that is compressed and dynamic. These methods use a 64-bit hash table to provide a basic de Bruijn graph implementation that can be updated.

### Other Related Compressed Data Structures

Some related compressed data structures are SeqOthello (Yu *et al*., 2018), SBT (Solomon and Kingsford, 2016), Split-SBT (Solomon and Kingsford, 2018), and Allsome-SBT (Sun *et al*., 2017). These methods index all *k*-mers or variants of *k*-mer indexes but do not provide graph information. For this reason, they are frequently applied to querying large collections of RNA-seq data but not genome assembly. Another indexing method is BIGSI (Bradley *et al*., 2017), which provides graph and can be viewed as a probabilistic colored de Bruijn graph.

Lastly, two other approaches are worthy of note because they merge the BWT of a set of strings. BWT-Merge (Sirén, 2016) is related to our work since the data structure we construct and store is similar to BWT. BWT-Merge merges two strings stored using BWT by using a reverse trie of one BWT to generate queries that are then located in the other BWT using FM-Index backward search. The reverse trie allows the common suffixes across multiple merge elements to share the results of a single backward search step. Thus, BWT-Merge finds the final rank of each full suffix completely, one suffix at a time. Finally, MSBWT (Holt and McMillan, 2014) is a method which merges the BWTs of multiple strings in a method similar to our own except applied to strings instead of graphs.

In Section 5, we compared the performance of VariMerge against Vari (Muggli *et al*., 2017)/Rainbowfish (Almodaresi *et al*., 2017), Bloom Filter Trie (Holley *et al*., 2016), Multi-BRWT (Karasikov *et al*., 2019) and Mantis (Pandey *et al*., 2018) / the method (Almodaresi *et al*., 2019). Among all mentioned methods in this section we chose these tools based on the following criteria: One method should be both graph-based and non-probabilistic (exact) for a fair comparison. Moreover, the authors of Mantis demonstrated that it (and therefore the recent work of Almodaresi *et al*. (2019)) are more efficient with respect to disk, memory and time than probabilistic method Split-SBT which itself is the efficient extension of SBT.

## 3 Preliminaries

As previously mentioned, Vari (Muggli *et al*., 2017) represents of the colored de Bruijn graph using BWT, and VariMerge, efficiently merges de Bruijn graphs that are represented in this manner. Here, we first define some basic notation and definitions concerning BWT, then we show how the de Bruijn graph and colored de Bruijn graph can be stored using BWT.

### 3.1 Basic Definitions and Terminology

Here, we begin with some basic definitions related to our representation. Throughout we consider a string X = X[1..*n*] = X[1]X[2] … X[*n*] of |X| = *n* symbols drawn from the alphabet [0..*σ* − 1]. For *i* = 1,…, *n* we write X[*i*..*n*] to denote the *suffix* of X of length *n* − *i* + 1, that is X[*i*..*n*] = X[*i*]X[*i* + 1] … X[*n*]. Similarly, we write X[1..*i*] to denote the *prefix* of X of length *i*. X[*i*..*j*] is the *substring* X[*i*]X[*i* + 1] … X[*j*] of X that starts at position *i* and ends at *j*.

#### Suffix arrays and suffix array intervals

The suffix array (Manber and Myers, 1993) SA_X_ (we drop subscripts when they are clear from the context) of a string X is an array SA[1..*n*] which contains a permutation of the integers [1..*n*] such that X[SA[1]..*n*] ≺ X[SA[2]..*n*] ≺ ⋯ ≺ X[SA[*n*]..*n*]. In other words, SA[*j*] = *i* if and only if X[*i*..*n*] is the *j*^th^ suffix of X in lexicographical order. Here, ≺ denotes lexicographic precedence.

For a string Y, the Y-interval in the suffix array SA_X_ is the interval SA[*s*..*e*] that contains all suffixes having Y as a prefix. The Y-interval is a representation of the occurrences of Y in X. For a character *c* and a string Y, the computation of *c*Y-interval from Y-interval is called a *left extension*.

#### BWT

Next, for a string Y, let F be the list of Y’s characters sorted lexicographically by the suffixes starting at those characters, and L be the list of Y’s characters sorted lexicographically by the suffixes starting immediately after those characters. (The names F and L are standard for these lists.) If Y[*i*] is in position *p* in F then Y[*i* − 1] is in position *p* in L. Moreover, if Y[*i*] = Y[*j*] then Y[*i*] and Y[*j*] have the same relative order in both lists; otherwise, their relative order in F is the same as their lexicographic order. This means that if Y[*i*] is in position *p* in L then (assuming arrays are indexed from 0) in F it is in position

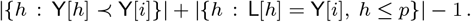

Finally, notice that the last character in Y always appears first in L. It follows that we can recover Y from L, which is the famous *Burrows-Wheeler Transform (BWT)* (Burrows and Wheeler, 1994) of Y.

#### FM-index and backward seach

Ferragina and Manzini Ferragina and Manzini (2005) first realized BWT can be used for indexing in addition to compression. Hence, if we know the range BWT(Y)[*i*..*j*] occupied by characters immediately preceding occurrences of a pattern *P* in Y, then we can compute the range BWT(Y)[*i*′..*j*′] occupied by characters immediately preceding occurrences of *cP* in Y, for any character *c*, since

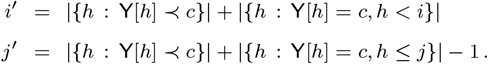

Notice *j*′ − *i*′ + 1 is the number of occurrences of *cP* in *S*. The essential components of an FM-index for Y are: (1) an array storing |{*h*:Y[*h*] ≺ *c*}| for each character *c* and, (2) a *rank* data structure for BWT(Y) that quickly tells us how often any given character occurs up to any given position. To be able to locate the occurrences of patterns in Y (in addition to just counting them), we can use a sampled suffix array of Y and a bitvector indicating the positions in BWT(Y) of the characters preceding the sampled suffixes.

Hence, we define the function rank(Y, *c, i*), for string Y, symbol *c*, and integer *i*, as the number of occurrences of *c* in Y[1..*i*]. Rank is used in *backward search* (Ferragina and Manzini, 2005) in order to compute left extension of a given string, i.e., the previous character.

### 3.2 Storage of de Bruijn Graph using BWT

Given a de Bruijn graph *G* = (*V, E*), we refer to the label of an edge *e* ∈ *E* as the *k*-mer corresponding to it, and denote it as label(*e*). Further, given *V*, we define the co-lexicographic (colex) ordering of *V* as the lexicographic order of their reversed labels ((*k* − 1)-mers).

We let F be the edges in *E* in colex order by their ending nodes, where ties are broken by their starting nodes, and let L be the edges in *E* sorted colex by their starting nodes, with ties broken by their ending nodes. We refer to the ordering of L as *Vari-sorted*. If we are given two edges *e* and *e*′ that have the same label then we are guaranteed that they have the same relative order in both F and L; otherwise, their relative order in F is the same as their labels’ lexicographic order. This means that if *e* is in position *p* in L, then in F it is in position

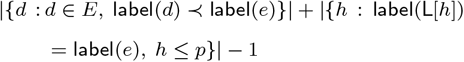

where ≺ denotes lexicographic precedence. We define the edge-BWT (EBWT) of *G* to be the sequence of edge labels sorted according to the ordering of the edges in L, so label(L[*h*]) = EBWT(*G*)[*h*] for all *h*. Therefore, if we have an array *D* storing |{*d*:*d* ∈ *E*, label (*d*) ≺ *c*}| for each character *c* and a fast rank data structure on EBWT(*G*) then given an edge’s position in L, we can quickly compute its position in F.

We let B_*F*_ be the bit vector with a 1 marking the position in F of the last incoming edge of each node, and let B_*L*_ be the bit vector with a 1 marking the position in L of the last outgoing edge of each node. Given a character *c* and the colex rank of a node *v*, we can use B_*L*_ to find the interval in L containing *v*’s outgoing edges. We can then search EBWT(*G*) to find the position of each outgoing edge^3^. Similarly, we can make similar queries about the incoming edges of a node *v* in an efficient manner using B_*F*_.

Therefore, briefly we explained how we can construct and represent a de Bruijn graph *G* = (*V, E*) with EBWT, B_*L*_, and B_*F*_ in a manner that allows for efficient navigation of the graph. An example of this representation is shown in Figure 1. Given this representation we can traverse the graph and recover incoming and outgoing edges. Next, we demonstrate how the labels (*k*-mers) can be recovered using this data structure.

**Fig. 1:**
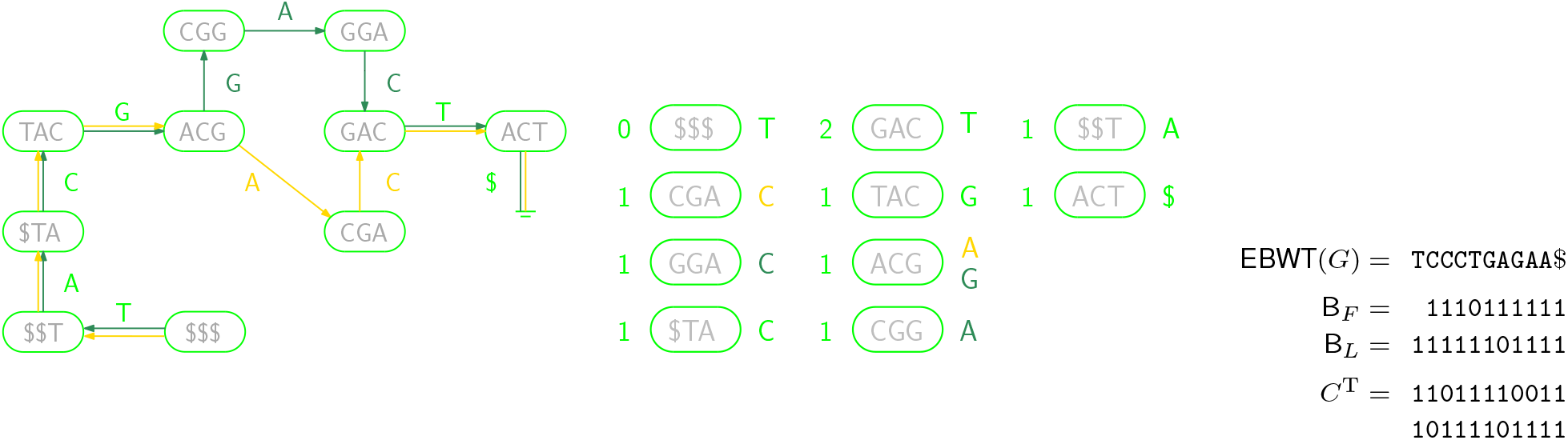
**Left:** A colored de Bruijn graph consisting of two individual graphs, whose edges are shown in yellow and green. All nodes to be present in both graphs are shown in lime. **Center:** The co-lexicographically sorted nodes, with the number of incoming edges shown on their left and the labels of the outgoing edges shown on their right. The edge labels are shown in yellow or green if the edges occur only in the respective graph, or lime if they occur in both. **Right:** The Vari representation of the colored de Bruijn graph: the edge-BWT is the list of edge labels. B_*F*_ and B_*L*_ are bit vectors to track the incoming and outgoing node degree, respectively. Finally, the binary array *C* (shown transposed) indicates which edges are present in which individual graphs.

#### Label recovery

We note that an important aspect of this succinct representation of the graph is that the (*k* − 1)-mers (nodes) and *k*-mers (edges) of the de Bruijn graph *G* are not explicitly stored in the above representation—rather they than can be *computed* (or recovered) from this representation. As previously mentioned, we can traverse the graph in a forward or reverse manner and recover incoming and outgoing edges of a given node *v*. Given this efficient traversal, we can recover the label of *v* by traversing the graph in a backward direction starting from *v*; given the label of *v* is a (*k* − 1)-mer we traverse backward *k* − 1 times. Therefore, we must add extra nodes and edges to the graph to ensure there is a directed path of length at least *k* − 1 to each original node. More formally, we augment the graph so that each new node’s label is a (*k* − 1)-mer that is prefixed by one or more copies of a special symbol $ not in the alphabet and lexicographically strictly less than all others. When new nodes are added, we are assured that the node labeled $^*k*−1^ is always first in colex order and has no incoming edges. Lastly, we augment the graph in a similar manner by adding an extra outgoing edge, labeled $, to each node with no outgoing edge. These “dummy nodes” are shown in Figure 1.

### 3.3 Storage of Colors

Given a multiset 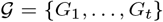 of individual de Bruijn graphs, we set *G* to be the union of those individual graphs and build the previously described representation for *G*. We also build and store a two-dimensional binary array C in which C[*i, j*] indicates whether the *i*th edge in *G* is present in the *j*th individual de Bruijn graph (i.e., whether that edge has the *j*th color). Hence, we store a given de Bruijn graph using EBWT, the described bit vectors, and a compressed color matrix. The combination of the above storage of the graph *G* plus the compressed matrix is the succinct storage of the graph.

## 4 Method

In this section, we first describe a naive merge procedure to motivate the use of the succinct merge algorithm; and then give the succinct merge algorithm in detail. We describe how to merge two colored de Bruijn graphs in both algorithms but note that it generalizes to an arbitrary number of graphs. Hence, we assume that we have two de Bruijn graphs *G*_1_ = (*V*_1_, *E*_1_) and *G*_2_ = (*V*_2_, *E*_2_) as input, which are stored as EBWT(*G*)_1_, B_*L*1_, B_*F*1_ and C_1_ and EBWT(*G*)_2_, B_*L*2_, B_*F*2_ and C_2_, respectively. We will output the merged graph *G_M_* = (*V_M_, E_M_*) stored in the same format as the input, more descriptively: a set of abbreviated edge labels EBWT(*G*)_*M*_, a bit vector that delimits their common origins B_*LM*_, the array B_*FM*_, and the color matrix C_*M*_.

### 4.1 A Naive Merge Algorithm

We recall from Section 3 that Vari does not store the edge labels (*k*-mers) of *G*–rather, they have to be computed from the succinct representation. We denote the edge labels for *G*_1_, *G*_2_, and *G_M_* as L_1_, L_2_, and L_*M*_, respectively. For example, if we want to reconstruct the *k*-mer AGAGAGTTA contained in *G*_1_ which is stored as `A` in EBWT(*G*)_1_, we need to backward navigate in *G*_1_ from the edge labeled `A` through *k* − 1 predecessor edges (`T, T, G`,…). We concatenate the abbreviated edge labels encountered during this backward navigation in reverse order to construct the label AGAGAGTTA. Thus, we could naively merge *G*_1_ and *G*_2_ by reconstructing L_1_ and L_2_, merging them into L_*M*_ and computing the succinct representation of L_*M*_, i.e., EBWT(*G*)_*M*_. We note that this algorithm requires explicitly building L_1_, L_2_ and L_*M*_ and thus, has a significant memory footprint.

### 4.2 The Succinct Merge Algorithm

We are now ready to describe the merge algorithm used as a component of VariMerge. The trick of the merge algorithm is to build the succinct data structure for *G_M_* without constructing L_1_ and L_2_, which will in turn reduce memory costs enormously.

#### 4.2.1 Intuitive Explanation of Succinct Merging

Before we give a detailed explanation of our algorithm we take a step back–abstract away the complexities of the succinct de Bruijn graph–and consider the simpler problem of merging two sorted lists of strings with the constraint that we can only examine a single character from each string at a time. We can solve this problem with a divide and conquer approach. First, we group all the strings in each list by their first character. This partially solves the problem, as we know all the strings in the first group from each list must occur in the output before all the strings in the second group in each list and so on. Thus, the problem is now reduced to merging the strings in the first group, followed by merging the strings in the second group and so on. Each of these merges can be addressed by again grouping the elements (i.e. subgroups of the initial groups) by examining the second character of each string. We can apply this step recursively until all characters of each string have been examined.

We draw the reader’s attention to the fact that our succinct colored de Bruijn graph representation is a space-efficient representation of the list of sorted *k*-mers (and (*k* − 1)-mers). Thus, we can apply this general algorithm but alter it in the following ways: 1) the nested grouping of the strings is rather a flat partitioning of the two lists into intervals, which is updated each time a character is processed, and 2) the actual merging is reserved for the end once all characters have been processed and their needed information accumulated into set of partitions of each list.

#### 4.2.2 Overview of the Algorithm

Now we return to the problem of merging succinct colored de Bruijn graphs. We refer to EBWT(*G*)_*M*_ and C_*M*_ as the *primary components* of the data structure and B_*FM*_ and B_*LM*_ as *secondary components*. We describe how to merge the primary components, and leave the details of how to merge the secondary components to the Supplement. The algorithm consists of two steps: (1) a planning step which plans the merge, and (2) a final execution step which executes the planned merge. In the planning step, we output a list of non-overlapping intervals for L_1_ andonefor L_2_. We refer to these lists as a *merge plan*, which is then used to execute the merge. There are *k* iterations of the planning algorithm (where *k* corresponds to the *k*-mer value). At each iteration of the algorithm a single character of the edge labels (*k*-mers) is processed, and the merge plan is revised (see Figure 3 in Supplement). After *k* iterations, we execute the merge plan.

#### 4.2.3 The Planning Step

We denote the merge plan as 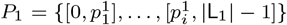, where each 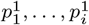 is an index in L_1_, and 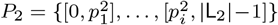, where each 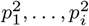 is an index in L_2_. We first initialize *P*_1_ and *P*_2_ to be single intervals covering L_1_ and L_2_, respectively (e.g. *P*_1_ = {[0, |L_1_| − 1]} and *P*_2_ = {[0, |L_2_| − 1]})(e.g.red merge plan in Figure 3). Next, we revise *P*_1_ and *P*_2_ in an iterative manner. In particular, we perform *k* consecutive revisions of *P*_1_ and *P*_2_, where *k* is the *k*-mer value used to construct *G*_1_ and *G*_2_—each revision of *P*_1_ and *P*_2_ is based on the next character^4^ of each edge label in L_1_ and L_2_ (e.g. green and blue merge plans in Figure 3). Thus, in order to fully describe the planning stage, we define (1) how the characters of the edge labels are computed and (2) how *P*_1_ and *P*_2_ are revised based on these characters.

##### Computing the next character of L_1_ and L_2_

We let *i* denote the current iteration of our revision of *P*_1_ and *P*_2_, where 1 ≤ *i* < *k*. We compute the next characters of L_1_ and L_2_ using two temporary character vectors 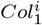 and 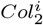, which are of length |L_1_| and |L_2_|, respectively. Conceptually, we define these vectors as follows: 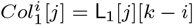 if *j* < *k* and otherwise 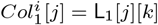, and 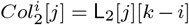 if *j* < *k*; and otherwise 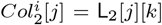. Since we do not explicitly build or store L_1_ and L_2_, we must compute 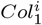 and 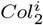. (Figure 4 shows how we calculate *Col^i^* based on *Col*^*i*−1^ at each iteration.)

##### Revising *P*_1_ and *P*_2_

We revise *P*_1_ and *P*_2_ based on 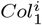 and 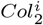 at iteration *i* by considering each pair of intervals in *P*_1_ and *P*_2_, i.e., *P*_1_ [*n*] and *P*_2_ [*n*] for *n* = 1,…, |*P*_1_|, and partitioning each interval into at most five (number of alphabets) sub-intervals. We store the list of sub-intervals of *P*_1_ and *P*_2_ as *SubP*_1_ and *SubP*_2_. Intuitively, we create *SubP*_1_ and *SubP*_2_ in order to divide *P*_1_ [*n*] and *P*_2_ [*n*] based on the runs of covered characters in 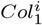 and 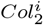—e.g., for each run of `A, C, G, T` or `$`. Next, we formally define this computation.

**Fig. 2:**
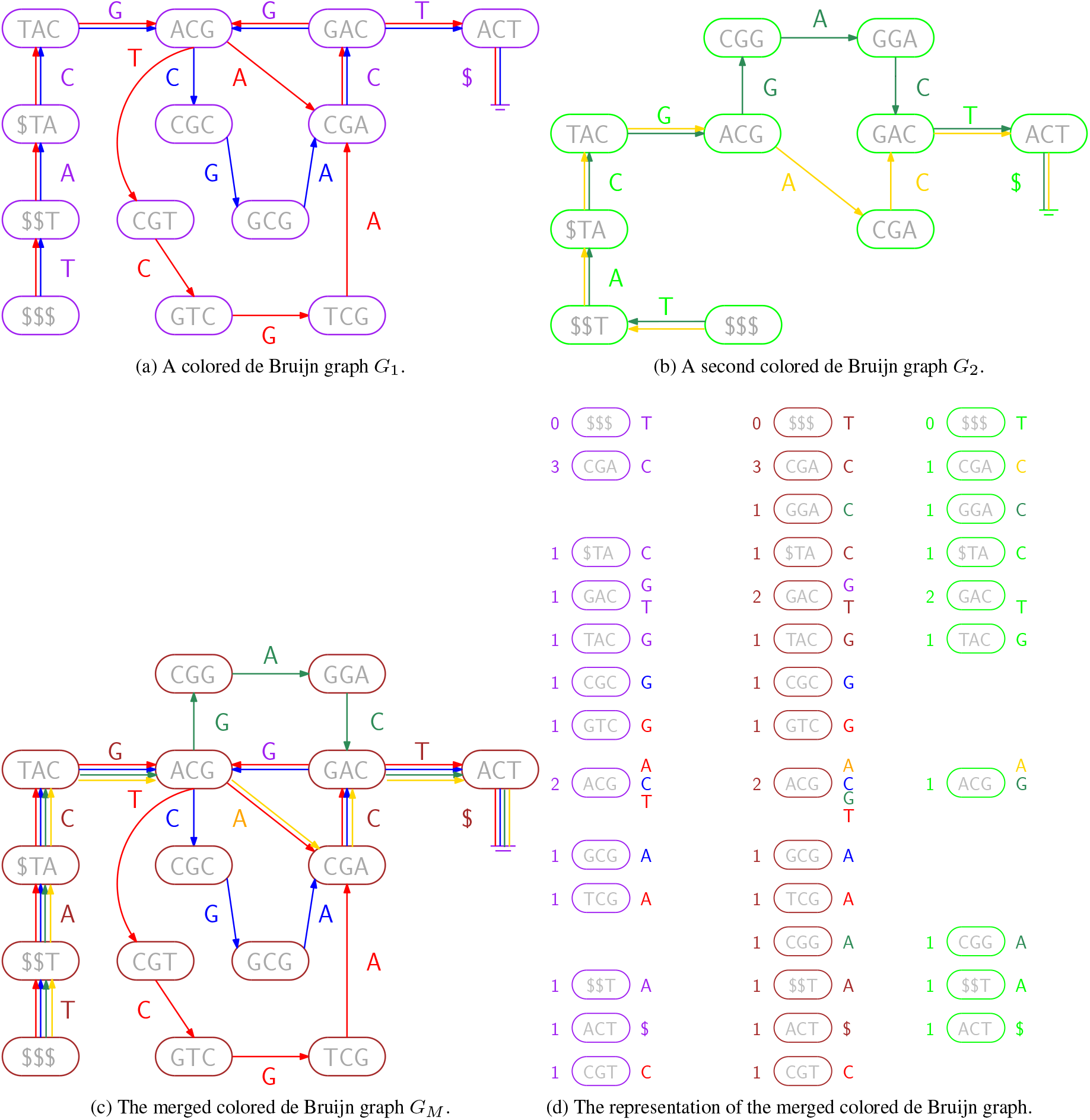
(**a**): A colored de Bruijn graph consisting of two individual graphs, whose edges are shown in red and blue. (We can consider all nodes to be present in both graphs, so they are shown in purple.) (**b**): A second colored de Bruijn graph, whose edges are green and yellow and lime represents presence in both graphs. (**c**): A colored de Bruijn graph merged from the two colored de Bruijn graphs. (**d**): The nodes for all three graphs arranged in columns (red and blue, merged, green and yellow). Each column is sorted into co-lexicographic order, with each node’s number of incoming edges shown on its left and the labels of its outgoing edges shown on its right. Vertical alignment illustrates how the merged components (center) are copied from either the left, the right, or both.

Thus, we partition *P*_1_ by first computing the subvector of 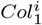 that is covered by *P*_1_[*n*], which we denote as 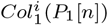, and computing the subvector of 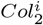 that is covered by *P*_2_[*n*], which we denote as 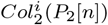. Next, given a character *c* in {$, `A, C, G, T`}, we populate *SubP*_1_[*c*] and *SubP*_2_[*c*] based on 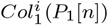 *and* 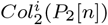 as follows: (1) we check whether *c* exists in either 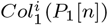 or 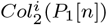; (2) if so, we add an interval to *SubP*_1_[*c*] covering the contiguous range of *c* in 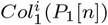 (or add an empty interval if 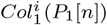 lacks any instances of *c*), and add an interval to *SubP*_2_[*c*] covering the contiguous range of *c* in 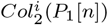 (or, likewise, add an empty interval if 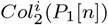 lacks any instances of *c*)^5^. Finally, we concatenate all the lists in *SubP*_1_ and *SubP*_2_ to form the revised plan 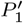 and 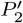. This revised plan 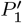 and 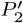 becomes the input *P*_1_ and *P*_2_ for the next refinement step. (Figure 3 shows three merged plans, including two refined ones (green and blue)).

**Fig. 3:**
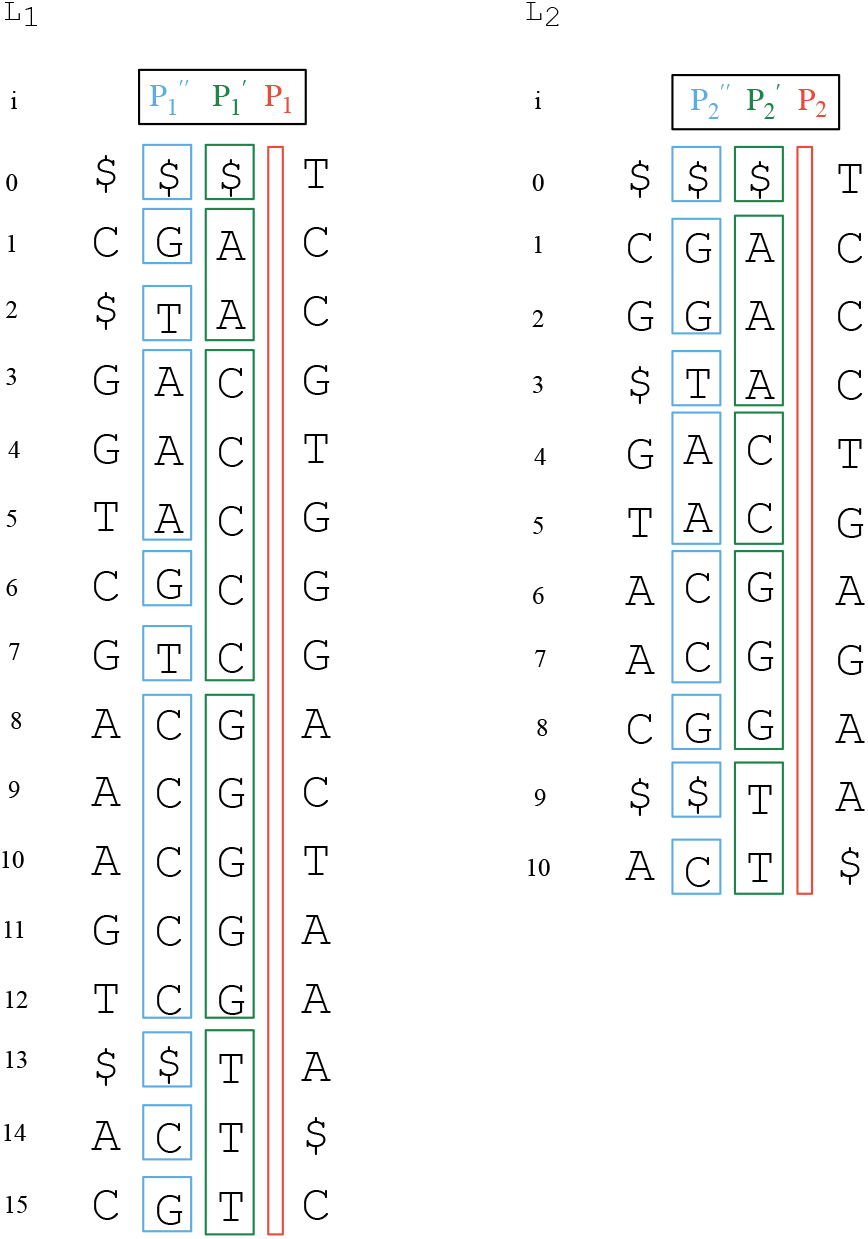
Three merge plans. The initial merge plan in red, where all elements of L_1_ and L_2_ are in the same equivalence class because no elements of either have been considered, e.g. *P*_1_ = {[0, 15]} and *P*_2_ = {[0, 10]}. Green is the first refinement, where a single character from each element in L_1_ and L_2_ are considered, e.g. 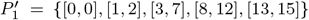 and 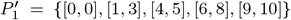. Blue is the third merge plan (second refinement), e.g. 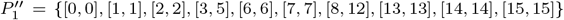 and 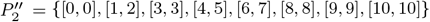.

We crafted the method above to maintain the property described in the following observation.

Observation 1. *Let P*_1_ *be a (partial) merge plan, and* 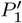 *its refinement by our merge algorithm, where ℓ*_1_,…, *ℓ_n_ are the elements in* L_1_ *that are covered by interval p_i_* ∈ *P*_1_ *and m*_1_,…, *m_o_ are the elements of* L_2_ *covered by interval q_j_* ∈ *P*_2_. *The following conditions hold: (1)* |*P*_1_| = |*P*_2_| *and* 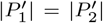; *(2) given any pair of elements where ℓ_a_* ∈ *p_i_, ℓ_b_* ∈ *p_j_ and* 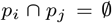 *there exists intervals* 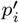 *and* 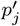 *in* 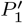 *such that* 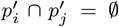 *and* 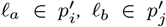; *and lastly, (3) given an interval p_i_ in P*_1_ *and the subsets of the alphabet used σ*_1_ ∈ *ℓ*_1_,…, *ℓ_n_ and σ*_2_ ∈ *m*_1_,…, *m_o_, then p_i_ will be partitioned into* |*SubP*_1_| = |*σ*_1_ ∪ *σ*_2_| *subintervals in* 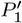.

We defined this observation for *P*_1_ but note that an analogous observation exists for *P*_2_.

#### 4.2.4 The Execution Step

We execute the merge plan by combining the elements of EBWT(*G*)_1_ that are covered by an interval in *P*_1_ with the elements of EBWT(*G*)_2_ that are covered by the equal position interval in *P*_2_ into a single element in EBWT(*G*)_*M*_. We note that when all characters of each label in L_1_ and L_2_ have been computed and accounted for, each interval in *P*_1_ and *P*_2_ will cover either 0 or 1 element of L_1_ and L_2_ and the number of intervals in *P*_1_ (equivalently *P*_2_) will be equal to |EBWT(*G*)_*M*_|. Thus, we consider and merge each pair of intervals of *P*_1_ and *P*_2_ in an iterative manner. We let 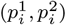 as the *i*-th pair of intervals. We concatenate the next character of EBWT (*G*)_1_ on to the end of EBWT(*G*)_*M*_ if 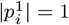. If 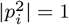 then we dismiss the next character of EBWT(*G*)_2_ since it is an abbreviated form of an identical edge to that just added. Next, if 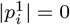 and 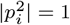, we copy the next character from EBWT(*G*)_2_ onto the end of EBWT(*G*)_*M*_.

We merge the color matrices in an identical manner by copying elements of C_1_ and C_2_ to C_*M*_. Again, we iterate through the plan by considering each pair of intervals. If 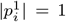 and 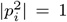 then we concatenate the corresponding rows of C_1_ and C_2_ to form a new row that is added to C_*M*_. If only one of 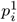 or 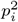 is non-zero then the corresponding row of C_1_ or C_2_ is copied to C_*M*_ with the other elements of the new row set to 0. Figure 3 in Supplement shows a complete illustration of VariMerge resulting colored de Bruijn graph.

The following theorem demonstrates the efficiency of our approach. The proof of the theorem of is in the Supplement.

##### Theorem 1.

*Given two de Bruijn graphs G*_1_ = (*V*_1_, *E*_2_) *and G*_2_ = (*V*_2_, *E*_2_) *constructed with integral value k such that, without loss of generality*, |*E*_1_| ≥ |*E*_2_|, *it follows that our merge algorithm constructs the merged de Bruijn graph G_M_ in O*(*m* · max(*k, t*))-*time, where t is the number of colors (columns) in* C_*M*_ *and m* = |*E*_1_|.

## 5 Results

In this section, we show the correctness of our method, the reduction in the resources used to build a colored de Bruijn graph by merging, and the comparison between state-of-the-art method for building exact colored de Bruijn graphs. We ran all performance experiments on a machine with two Xeon E5-2640 v4 chips, each having 10 2.4 GHz cores. The system contains 755 GB of RAM and two ZFS RAID pools of 9 disk each for storage. We report wall clock time and maximum resident set size from Linux.

### 5.1 Validation on E coli

In order to validate the correctness of our approach, we generated two succinct colored de Bruijn graphs with two sets of three *E. coli* assemblies each, merged them, and verified its equivalence to a six color graph built from scratch using Vari. We obtained the six sub-strains of *E. coli* K-12 from NCBI. Each of the genomes contained approximately 4.6 million base pairs and had a median GC content of 49.9%. This experiment tests that the merged colored de Bruijn graph built by VariMerge is equivalent to that produced by building the graph without merging (i.e., with Vari alone). We ran a bubble calling algorithm on both six colored graphs and found the identical set of bubbles between them. Further, we found VariMerge produced files on disk that were bit-for-bit identical to those generated by Vari, demonstrating they construct equivalent graphs and data structures. We leave the details of this validation to the supplement (see Table 4 in the supplement).

### 5.2 Demonstration of Large-scale Construction and Incremental Updates

We downloaded the sequence data from for 16,000 Salmonella strains (NCBI BioProject PRJNA18384) and divided them into four sets of 4,000 strains, which we label 4A, 4B, 4C, and 4D. From these datasets we construct a colored de Bruijn graph for 8,000 strains, to demonstrate the efficiency of the merge process, and 16,000 strains, to demonstrate scalability. The exact accessions for each dataset is available in our repository.

In order to measure the effectiveness of VariMerge for the proposed divide-and-conquer method of building large graphs, we constructed the colored de Bruijn graph using Vari for a set of 4,000 salmonella assemblies (4A). This took 8 hours 46 minutes, 1 TB of external memory, and 136 GB of RAM to build the graph for 4,000 strains. We then built a graph for a second set of 4,000 assemblies (4B) using 10 hours 40 minutes, 1.5 TB of external memory, and 137 GB of RAM. We merged these two 4,000 sample graphs (i.e. 8AB) using our proposed algorithm in 2 hours 1 minutes, no external memory, and 10 GB of RAM. “’0” is shown in the external memory column of Table 1 for merge since no external memory is ever used for merging. Thus the VariMerge method required a combined 137 GB of RAM, 26 hours 30 minutes of runtime to produce the graph for 8,000 strains. We denote this graph with 8,000 strains as 8AB. In contrast, running Vari on the same 8,000 strains required 37 hours 27 minutes, 4.6 TB of external memory and 271 GB of RAM. Thus VariMerge reduced runtime by 11 hours, reducing RAM requirements to 134 GB, and reducing external memory requirements by 3.1 TB.

**Table 1.** Breakdown of the memory, disk and time usage of VariMerge to build the colored de Bruijn graph for 8,000 strains. The VariMerge method consists of running Vari on subsets of the population (4A and 4B) and then merging the results with our proposed merge algorithm (denoted Merge here). We list the resources used for both individual runs of Vari, the Merge required, and the combined resources. The combined resources consists of the total time and maximum space used across all three components of VariMerge used in this dataset. No external memory is needed for merging itself so “0” is in the external memory column for Merge

| Program and Dataset | Input Stats |  | de Bruijn Graph |  |  | Color Matrix |  |  | Combined Requirements |  |  |  |
| --- | --- | --- | --- | --- | --- | --- | --- | --- | --- | --- | --- | --- |
|  | *k*-mers | Colors | RAM | Time | Size | RAM | Time | Size | RAM | Ext. Mem. | Time | Size |
| Vari(4A) | 1.1 B | 4,000 | 136 GB | 8 h 46 m | 0.31 GB | 52 GB | 1 h 39 m | 51.2 GB | 136 GB | 1 TB | 10 h 25 m | 51 GB |
| Vari(4B) | 1.5 B | 4,000 | 137 GB | 10 h 40 m | 0.52 GB | 54 GB | 2 h 22 m | 52.5 GB | 137 GB | 1.5 TB | 13 h 2 m | 53 GB |
| Merge(4A, 4B) | 2.4 B | 8,000 | 10 GB | 2 h 1 m | 0.63 GB | 117 GB | 1 h 2 m | 106 GB | 117 GB | 0 TB | 3 h 3 m | 106 GB |
| VariMerge | 2.4 | 8,000 | 137 GB | 21 h 27 m | 0.63 GB | 117 GB | 5 h 3 m | 117 GB | 137 GB | 1.5 TB | 26 h 30 m | 106 GB |

We further used this facility to merge two more 4,000 color graphs (i.e. 4C + 4D) (Table 2). We denote the resulting graph as 8CD. We then merged this 8,000 sample graph with the aforementioned 8,000 graph to produce a succinct colored de Bruijn graph of 16,000 samples (i.e. 8AB + 8CD).

**Table 2.** Breakdown of the peak memory, peak disk and time required by VariMerge to build the colored de Bruijn graph for 16,000 strains of Salmonella. We note VariMerge includes the resources required of the two 4,000 runs of Vari (i.e. Vari(4A) and Vari(4B)) and the merge run (i.e. Merge(4A, 4B)) from Table 1. No extra external memory is needed for merging so “0” is in the external memory column for Merge

| Program and Dataset | Input Stats |  | de Bruijn Graph |  |  | Color Matrix |  |  | Combined Requirements |  |  |  |
| --- | --- | --- | --- | --- | --- | --- | --- | --- | --- | --- | --- | --- |
|  | *k*-mers | Colors | RAM | Time | Size | RAM | Time | Size | RAM | Ext. Mem. | Time | Size |
| Vari(4C) | 1.7 B | 4,000 | 135 GB | 10 h 53 m | 0.46 GB | 53 GB | 2 h 34 m | 51.8 GB | 135 GB | 1.6 TB | 13 h 27 m | 52 GB |
| Vari(4D) | 2.4 B | 4,000 | 137 GB | 14 h 35 m | 0.67 GB | 59 GB | 3 h 37 m | 57.9 GB | 137 GB | 2.34 TB | 18 h 12 m | 59 GB |
| Merge(4C, 4D) | 3.8 B | 8,000 | 17 GB | 2 h 59 m | 1.00 GB | 118 GB | 57 m | 107 GB | 118 GB | 0 TB | 3 h 56 m | 108 GB |
| Merge(8AB, 8CD) | 5.8 B | 16,000 | 25 GB | 4 h 53 m | 1.60 GB | 254 GB | 2 h 10 m | 232 GB | 254 GB | 0 TB | 7 h 3 m | 233 GB |
| VariMerge | 5.8 B | 16,000 | 137 GB | 54 h 47 m | 1.60 GB | 254 GB | 14 h 21 m | 232 GB | 254 GB | 2.34 TB | 69 h 8 m | 233 GB |

In order to measure the effectiveness of VariMerge for incremental additions to a graph that holds a growing population of genomes, we started with the colored de Bruijn graph of 16,000 salmonella assemblies. We then constructed a second graph for a singleton set of just one additional assembly. Next, we ran our proposed merge algorithm on these two graphs. VariMerge took 69 hours 8 minutes, 2.34 TB of external memory, and 254 GB of RAM to build the graph for 16,000 strains. To build a single colored de Bruijn graph for an additional strain, Vari took 7 seconds, 460 MB of external memory, and 2.3 GB of RAM. Our proposed merge algorithm took 7 hours 9 minutes, no external memory, and 254 GB of RAM to merge the 16,000 color graph with the 1 color graph. This is an order of magnitude faster than the almost 70 hours it would take to build the same 16,001 color graph from scratch.

### 5.3 Comparison to Existing Methods

We compare our method to the existing space- and memory-efficient colored de Bruijn graphs. To accomplish this, we ran Bloom Filter Trie Holley *et al*. (2016), Vari (Muggli *et al*., 2017) / Rainbowfish (Almodaresi *et al*., 2017), Mantis (Pandey *et al*., 2018) / the method of Almodaresi *et al*. (2019), and Multi-BRWT (Karasikov *et al*., 2019). The peak RAM, peak disk usage, and running time required by the methods to construct a colored de Bruijn graph for 4,000, 8,000, and 16,000 strains is shown in Table 3. All methods were ran with default parameters and *k* = 32, except from Bloom Filter Trie which can only work with *k* values which are multiples of 9, hence we ran it with *k* = 27. Also we ran Mantis on 4,000 and 8,000 strains with suggested log-slot value for large experiments which is 33, however we had to increase this value to 36 for running it on 16,000 strains due to “auto resize failed” error. All method are exact, colored de Bruijn graph construction methods.

**Table 3.**
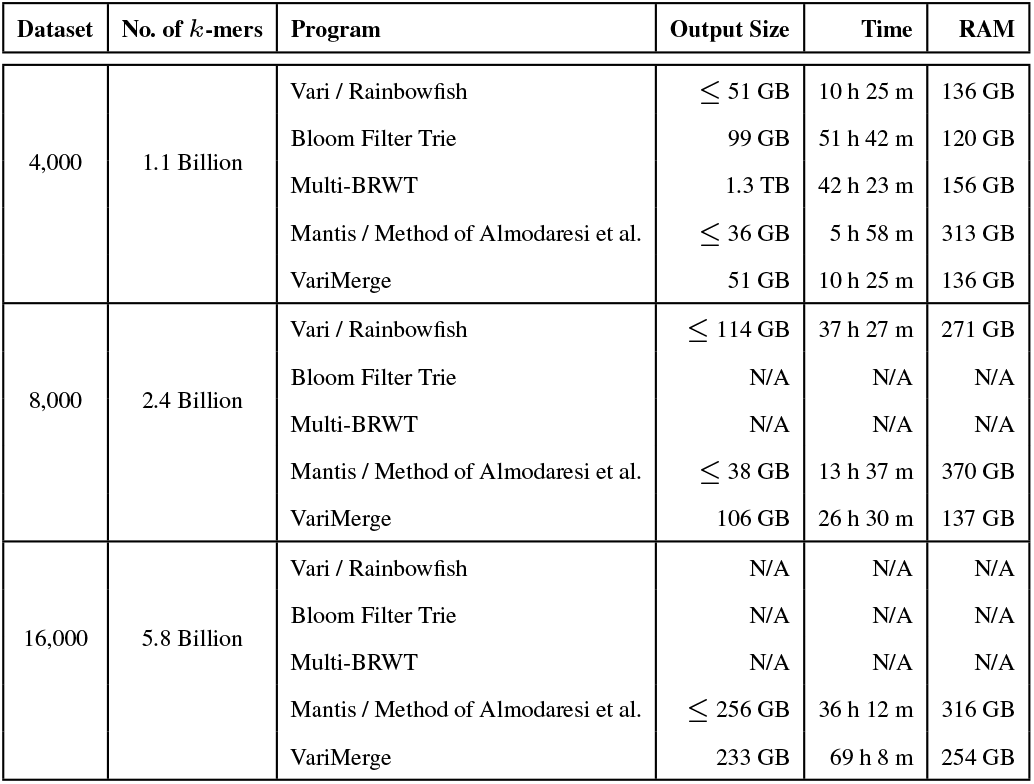
Comparison between building a space-efficient colored de Bruijn graph construction methods for 4,000, 8,000, and 16,000 Salmonella strains using VariMerge verses competing methods. We report N/A for any method that exceeded 72 CPU hours, 4 TB of disk space, and 750 GB of memory. ‘≤’ denotes the value for a base method that has a later an add-on method. We anticipate add-on methods to compress better but will still consume the resources shown for their base method because they reuse base the method’s output. We measured RAM as Max Resident Set Size. Mantis authors noted their use of memory mapped I/O means this reveals opportunitic consumption and not necessarily requirement for their program.

Since Rainbowfish first runs Vari and then compresses, the peak RAM, peak disk usage, and time will be at least that of Vari for construction; this is why they are shown together in Table 3. Similarly, we report peak RAM, peak disk usage and time of Mantis together with that of the method of Almodaresi *et al*. (2019) because the latter method first runs Mantis and then compresses the output of Mantis. These points are also discussed in the related work. Again, because we are interested in construction (not exclusively compression) we are interested in peak RAM and external memory usage.

We report N/A in Table 3 for any method that exceeded 72 hours, and/or our RAM and external memory quota. We had 4 TB of disk space for each method (20 TB in total) and 750 GB of RAM. All methods performed reasonably well on 4,000 strains. Bloom Filter Trie required the most time, and Multi-BRWT required the most space. This was unsurprising as Multi-BRWT focuses on compression of the color matrix and not graph (see Related Work). Mantis (and it’s successor (Almodaresi *et al*., 2019)) required the largest peak memory (313 GB) but had the lowest disk usage (35 GB). With 4,000 strains all the methods were capable of running given the constraints. With 8,000 strains Vari and Rainbowfish required more than 4 TB of disk space, and both Bloom Filter Trie and Multi-BRWT required more than 72 hours. Mantis and VariMerge were the only ones capable of building the colored de Bruijn graph with 8,000 strains. Mantis was more efficient with respect to final output and VariMerge was more efficient with respect to RAM consumed. Lastly, only Mantis and VariMerge were capable of building the colored de Bruijn graph with 16,000 strains. Yet, as shown in the previous section, VariMerge allows the graph to be updated with new data via merging. Mantis (as well as Vari and Rainbowfish) cannot update the graph without reconstruction.

Lastly, VariMerge has several practical advantages over competing methods. It allows large values of *k*, i.e., *k* ≤ 64, and allows for efficient queries of the form: given a color *c*, return all *k*-mers that have that color. Competing methods cannot perform such queries without the input files or cannot scale to large datasets. We discuss this more in the conclusions.

## 6 Conclusion

In this paper, we develop a method to build the colored de Bruijn graph by merging smaller graphs in a resource-efficient manner. This allows the colored de Bruijn graph to be constructed for large datasets, and provides an efficient means to update the colored de Bruijn graph. This latter point is particularly important as data is frequently added to public databases. We leave optimizing partition size as an area for future work.

Lastly, we mention that due to the underlying graph data structure of VariMerge has some advantages over competing methods. First, VariMerge allows arbitrary *k* up to 64 while other tools are more restricted; BFT requires *k* to be multiples of 9 and Mantis only supports up to *k*=32. Second, the node labels can be recovered from the index alone, allowing the following queries to be performed: given a particular color *c*, what *k*-mers have color *c*. These queries can be accomplished by VariMerge by scanning a particular column of the color matrix, and can be used for comparing samples, i.e., how many *k*-mers are shared between sample *x* and sample *y*. While the Mantis authors have suggested an approach to supporting this query, it is unimplemented and requires the input files, increasing the size of the data structure. See supplement for a full explanation.

## 7 Personal Communication with Authors of Mantis

On Thu, Jan 24, 2019 at 7:44 PM Rob Johnson wrote:

First of all, thank you for taking the time to have these discussions while under a deadline. I really appreciate it! I also appreciate the effort you put into running mantis under ulimit. It’s unfortunate that ulimit has no affect on recent linux versions.

But I have great confidence that the memory usage is not 313GB. Even though Mantis maps a lot of memory, it’s working set is much, much smaller than the total memory it maps. If there were any memory pressure on the system, the OS could reclaim that RAM and use it for something else, and Mantis would hardly be affected. We hope to get you a cgroups script that can properly limit mantis’ resident size.

As for returning all the kmers in the same sample that contains some kmer x, I don’t think that is the raison d’etre of a CDBG. For example, the query, “In which samples does this k-mer occur?” is very common. But nonetheless I believe this can be done trivially with Mantis as follows: 1. Keep all the squeakr files^6^. 2. Query in mantis for the sample containing x. This takes O(samples) time.

Then enumerate all the kmers from that sample’s squeakr file. This takes O(size of output) time, i.e. constant output complexity. This will increase disk usage (since you need to keep all the squeakrs), but I believe it would be competitive, in all regards, with everything in the table you sent before.

Best, Rob

## 8 Proof of Theorem 1

### Theorem 1.

Given two de Bruijn graphs *G*_1_ = (*V*_1_, *E*_2_) and *G*_2_ = (*V*_2_, *E*_2_) constructed with integral value *k* such that, without loss of generality, |*E*_1_| ≥ |*E*_2_|, it follows that our merge algorithm constructs the merged de Bruijn graph *G_M_* in *O*(*m* · max(*k, t*))-time, where *t* is the number of colors (columns) in C_*M*_ and *m* = |*E*_1_|.

### Proof.

In our merge algorithm, we will perform *k* refinements of *P*_1_ and *P*_2_ after they are initialized. We know by definition and Observation 1 that |*P*_1_| ≤ |L_1_|, *P*_2_ ≤ |L_2_|, 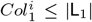 and 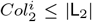 at each iteration *i* of the algorithm. Further, it follows from Observation 1 that a constant number of operations are performed to *P*_1_, *P*_2_, 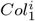 and 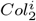. We populate C_*M*_ in the last step of merging the primary components of the data structure. Since the C_*M*_ is a bit matrix of size *k* by *t*, it follows that this step will take time *O*(*m* max(*k, t*))-time. Hence, if *k* ≤ *t* the merge algorithm will take *O*(*mk*)-time; otherwise it will take *O*(*mt*)-time (since populating C_*M*_ will dominate in this case).

## 9 Details Concerning Strains of E.coli K-12

Table 4 describes the accession number, sub-strain and genome length of the E.coli genomes used for validation of our construction and bubble-calling.

**Table 4.** Characteristics of the substrains of E. coli K-12 used to test the performance and accuracy of VariMerge

| Accession Number | Sub-strain | Genome Size |
| --- | --- | --- |
| AP009048 | W3110 | 4,646,332 bp |
| CP009789 | ER3413 | 4,558,660 bp |
| CP010441 | ER3445 | 4,607,634 bp |
| CP010442 | ER3466 | 4,660,432 bp |
| CP010445 | ER3435 | 4,682,086 bp |
| U00096 | MG1655 | 4,641,652 bp |

We validate VariMerge by generating two succinct colo

## 10 Merging the secondary components

### Delimiting common origin with B_*LM*_

We prepare to produce B_*LM*_ in the planning step by preserving a copy of the merge plan after *k* − 1 refinement iterations as *S*_*k*−1_. After *k* − 1 refinement steps, our plan will demarcate a pair of edge sets where their labels have identical *k* − 1 prefixes. Thus, whichever merged elements in EBWT(*G*)_*M*_ result from those demarcated edges will also share the same *k* − 1 prefix. Therefore, while executing the primary merge plan, we also consider the elements covered by *S*_*k*−1_ concurrently, advancing a pointer into EBWT(*G*)_1_ or EBWT(*G*)_2_ every time we merge elements from them. We form B_*LM*_ by appending a delimiting 1 to B_*LM*_ (again, indicating the final edge originating at a node) whenever both pointers reach the end of an equal rank pair of intervals in *S*_*k*−1_’s lists.

### Delimiting common destination with flags_*M*_

We produce flags in a similar fashion to B_*LM*_ but create a temporary copy of *S*_*k*−2_ in the planning stage after *k* − 2 refinement iterations instead of *k* − 1. In this cases, the demarcated edges are not strictly those that share the same destination; only those edges that are demarcated and share the same final symbol. Thus, in addition to keeping pointers into EBWT(*G*)_1_ or EBWT(*G*)_2_, we also maintain a vector of counters which contain the number of characters for each (final) symbol that have been emitted in the output. We reset all counters to 0 when a pair of delimiters in *S*_*k*−2_ is encountered. Then, when we append a symbol onto EBWT(*G*)_*M*_, we consult the counters to determine if it is the first edge in the demarcated range to end in that symbol. If so, we will not output a flag for the output symbol; otherwise, we will.

## 11 Illustration of merge plans

In this section we provide an example of merge plans in two conceptual edge lists L_1_ and L_2_ from graphs *G*_1_ and *G*_2_ in Figure 3, where *k* = 4. (Note that the edge labels are not stored explicitly; instead at each iteration the next letter is computed. See Figure 4). As mentioned before the merge plan is defined as 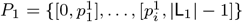 where each 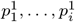 is an index in L_1_, and 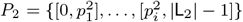, where each 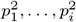 is an index in L_2_. Next, all revisions of *P*_1_ and *P*_2_ are computed iteratively. In the below example we start with the red merge plans *P*_1_ and *P*_2_ covering the whole L_1_ and L_2_ respectively. In the next iteration, based on the next letter (the order is right to left), we revise every interval into at most five subintervals (five being the number of alphabets: {$, `A, C, G, T`}). In this example see how *P*_1_ and *P*_2_ in red are revised to 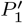 and 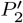 in green each with five intervals. Similarly every interval in 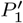 and 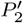 are revised to at most five subintervals making 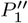 and 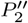. See the caption of Figure 3 for the intervals that each merge plan covers.

**Fig. 4:**
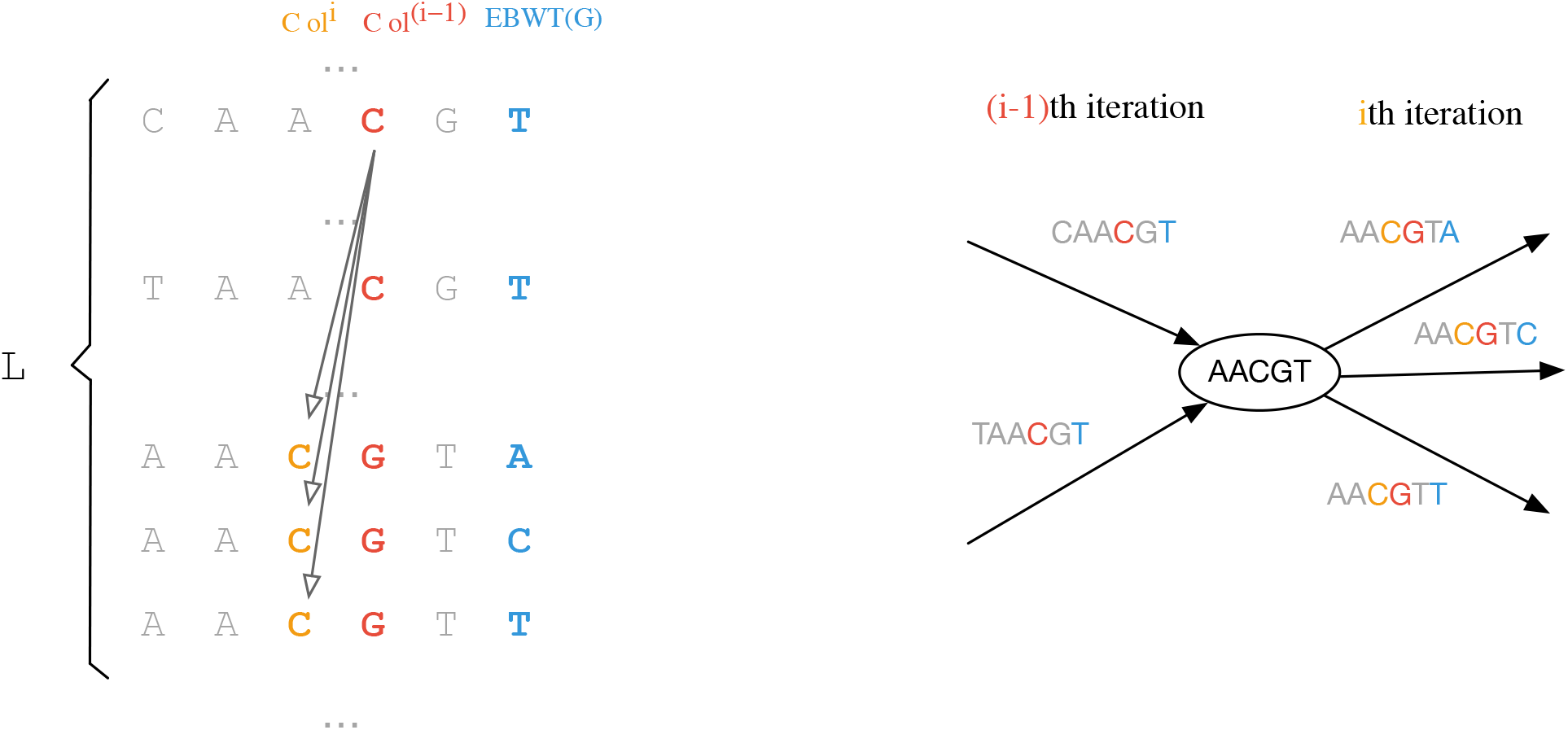
Method for populating *Col^i^* based on *Col*^*i*−1^ and graph navigation. Colored (blue, red and orange) nucleotides represent data that is in memory and valid. Grey represents data that is stored in external memory in Vari but is computed as needed and only exists ephemerally in VariMerge. Thus, only three columns are ever present in memory, which is a significant memory savings relative to the full set of edge labels. The three resident vectors are 1.) EBWT(*G*) (which is always present and used for navigation), 2.) *Col*^*i*−1^ which is already completely populated when a new column to the left, 3.) (*Col^i^*) is being generated. As one can see in the associated graph representation on right, the successor edges have almost the same label but shifted by one position. e.g. the red colored `C` in *i* − 1-th iteration in position 4 is shifted to position 3 and turned into orange in *i*-th iteration.

## 12 Illustration of computing the next character of L

In this section we provide an example of how we can navigate the set of edge labels without explicitely storing them. We compute the next character at each iteration with only three columns present in memory (shown with colors blue, red and orange). See the caption of Figure 4 for more details.

## Footnotes

1 Since Rainbowfish first constructs Vari and then compresses, its peak memory and disk usage is the same.

2 Since Almodaresi *et al*. (2019) constructs Mantis and then compresses, its peak space and disk usage is the same.

3 In practice, we incorporate the bits of *B_F_* as flags on EBWT(*G*) and use them to obtain the colex order of *v* but omit the discussion here for simplicity. We refer the reader to (Bowe *et al*., 2012) for a full discussion of this aspect and the supplement for our handling here.

4 We recall that FM-index stores the last character of each edge label and we do not have access to L_1_ and L_2_. Therefore, we are processing the characters of L_1_ and L_2_ from right to left. Thus, the “next” character is the preceding character of an edge label.

5 We are guaranteed by the definition of our data structure that any instances of *c* in 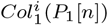 will be in a contiguous range, and likewise, any instances of *c* in 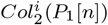 will also be in a contiguous range

6 Authors note: The size of the input files was 158 GB, 319 GB, and 656 GB for the 4,000, 8,000, and 16,000 strains considered in this paper.

